# Chromosome-length genome assemblies of cactophilic *Drosophila* illuminate links between structural and sequence evolution

**DOI:** 10.1101/2022.10.16.512445

**Authors:** Kyle M. Benowitz, Carson W. Allan, Coline C. Jaworski, Michael J. Sanderson, Fernando Diaz, Xingsen Chen, Luciano M. Matzkin

**Affiliations:** Department of Entomology, University of Arizona, Tucson, AZ, USA; Department of Biology, Austin Peay State University, Clarksville, TN, USA; Department of Zoology, Cambridge University, Cambridge, UK; Université Côte d’Azur, INRAE, CNRS, UMR ISA, 06000 Nice, France; Department of Ecology and Evolutionary Biology, University of Arizona, Tucson, AZ, USA; Department of Biology, Colgate University, Hamilton, NY, USA; BIO5 Institute, University of Arizona, Tucson, AZ, USA

## Abstract

A thorough understanding of adaptation and speciation requires model organisms with both a history of ecological and phenotypic study as well as a robust set of genomic resources. For decades, the cactophilic *Drosophila* species of the southwestern US and northern Mexico have fit this profile, serving as a crucial model system for understanding ecological adaptation, particularly in xeric environments, as well as the evolution of reproductive incompatibilities and speciation. Here, we take a major step towards gaining a complete molecular description of this system by assembling and annotating seven chromosome-length *de novo* genomes across the three species *D. mojavensis, D. arizonae*, and *D. navojoa*. Using this data, we present the most accurate reconstruction of the phylogenetic history of this clade to date. We further demonstrate a relationship between structural evolution and coding evolution both within and between species in this clade, and use this relationship to generate novel hypotheses for adaptation genes. All of our data are presented in a new public database (cactusflybase.arizona.edu), providing one of the most in-depth resources for the analysis of inter- and intraspecific evolutionary genomic data.

## Introduction

The fundamental goal of evolutionary genetics is to link phenotypic adaptation to genomic variation (Lewontin 1974). Importantly, the causality of this link, for practical purposes, can be viewed as bidirectional. It is essential to use genomic approaches to ascribe genetic underpinnings to previously identified adaptive phenotypes. Such top-down approaches are needed to answer fundamental questions regarding the type and number of genes underlying adaptation and the predictability of these processes, among others (Orr 2005; Barrett and Hoekstra 2011). On the other hand, it is equally as necessary to draw conclusions *a posteriori* from genomic comparisons to generate hypotheses about cryptic or otherwise understudied phenotypes that may be contributing to ecological adaptation and speciation (Benowitz et al. 2020). With this type of bottom-up approach, genomic data may be repurposed to benefit studies of organismal natural history (Holmes et al. 2016, Sherman et al. 2016).

What genomic data is precisely needed for these purposes? For much of the genomic era, understanding genome-wide variation specifically meant understanding variation at the level of the gene. Molecular evolution at the level of the gene remains, and always will remain, a fundamental aspect of evolutionary genomic practice. However, evidence is incontrovertible that structural chromosomal variants play essential roles in adaptation and speciation. Gene duplication has long been known to be a major driver of phenotypic adaptation (Ohno 1975), while large chromosomal inversions play fundamental roles in adaptation and speciation (Noor et al. 2001; Kirkpatrick and Barton 2006). More recently, though, it is increasingly recognized that a broader variety of genomic rearrangements, including sequence gain, loss and transposition via transposable elements (Casacuberta and González 2013; Schrader and Schmitz 2019), microinversions (Redmond et al. 2020; Connallon and Olito 2021), and chromosomal fusions (Wellband et al. 2019) may also contribute to adaptation. Smaller structural variants, including insertions, deletions, and transpositions have also been increasingly shown to be implicated in speciation (Zhang et al. 2021). Following this, efforts are ongoing to create reproducible approaches to identify all types of structural variation and quantifying their evolution across species and populations (Chakraborty et al. 2018; Wala et al. 2018; Goel et al. 2019; Heller and Vingron 2019; O’Donnell and Fischer 2020).

One challenge presented by the focus on more nuanced types of structural variation is that the fragmented, short-read assemblies that have been predominant in the world of non-model genomics may no longer suffice. Although these assemblies have been instrumental in facilitating gene expression studies and answering a wide variety of otherwise inaccessible questions regarding molecular evolution and the evolution of gene family content across a broad taxonomic range (Ellegren 2014), they offer an incomplete insight into gene duplication and none into the presence of larger structural variants (Chakraborty et al. 2018; Pollard et al. 2018; van Dijk et al. 2018). For this reason, the past several years have seen an increased emphasis on producing highly contiguous or chromosome-length genome assemblies for a broader range of organisms (Hotaling et al. 2021; Kim et al. 2021; Rhie et al. 2021). Fortunately, these efforts are being aided by both decreasing costs of long-read sequencing and the further development of methodologies to improve long-read assemblies (Amarasinghe et al. 2020; Jaworski et al. 2020; De Coster et al. 2021; Whibley et al. 2021).

Given the two-way street between genomic information and phenotypic and ecological information, we propose that the most promising study organisms will be those wherein hypotheses in both directions can be effectively leveraged; in other words, ecologically rich, tractable genomic systems with substantive empirical foundations in both areas. The cactophilic *Drosophila* within the *mulleri* complex of the *repleta* group neatly fit this description. Cactophilic flies have adapted to living in xeric environments by making a habitat of necrotic cactus tissue, where larvae develop and all life stages feed on yeasts (Fogleman et al. 1981; 1982) and bacteria (Fogleman and Foster 1982) proliferating in the necrosis, which is highly toxic (Kircher 1982; Fogleman and Heed 1989; Fogleman and Danielson 2001). As predicted, the transition to a cactophilic life-history has required adaptations to harsh environmental conditions including high temperatures (Schnebel and Grossfield 1984; Krebs 1999; Fasolo and Krebs 2004; MacLean et al. 2019; Shaible and Matzkin 2022), low humidity (Gibbs and Matzkin 2001; Matzkin et al. 2007; Matzkin et al. 2009), and high toxicity (Guillén et al. 2015).

In addition to the novel colonization of their habitat, there has also been extensive ecological divergence within the cactophilic group. One subclade that has received particular attention is the *Drosophila mojavensis* species cluster, consisting of the three species *D. mojavensis, D. arizonae*, and *D. navojoa* (Matzkin 2014). This clade, which has diversified within the last few million years (Russo et al. 1995; Matzkin and Eanes 2005; Reed et al. 2007, Smith et al. 2012), inhabits a range of cactus hosts and habitat types (Matzkin 2014). Within *D. mojavensis*, there are four geographically and genetically distinct populations that largely (but not exclusively) inhabit single, distinct host cacti (Matzkin 2014; Etges 2019): one in the Sonoran Desert inhabiting organ pipe cactus (*Stenocereus thurberi*), one in Baja California inhabiting agria (*Stenocereus gummosus*), one in the Mojave Desert inhabiting red barrel cactus (*Ferrocactus cylindraceus*), and one on Santa Catalina Island (CA) inhabiting prickly pear (*Opuntia littoralis*). Its sibling species, *D. arizonae*, is a generalist, inhabiting multiple cactus species within its range from Guatemala to southern California (Fellows and Heed 1972; Heed 1978; 1982). The outgroup, *D. navojoa* from central Mexico, is a specialist on prickly pear (*O. wilcoxii*; Heed 1982). These distinctions within and between species have proved to be an extremely fruitful substrate for hypotheses regarding phenotypic adaptation in many forms, including heat tolerance (Diaz et al. 2021a), desiccation resistance (Matzkin et al. 2007; Rajpurohit et al. 2013), chemical adaptation (Starmer et al. 1977); life-history (Etges 1990), olfaction (Date et al. 2013; Crowley-Gall 2016; 2019; Nemeth et al. 2018; Ammagarahalli et al. 2021), and behavior (Newby and Etges 1998; Coleman et al. 2018). Additionally, the recent divergence within and between species in the *D. mojavensis* species complex has made this clade into a model system for speciation and the evolution of reproductive incompatibilities (Markow 1981; Pantazidis and Zouros 1988; Zouros et al. 1988; Etges 1992; Knowles and Markow 2001; Miller et al. 2003; Pitnick et al. 2003; Reed and Markow 2004; Massie and Markow 2005; Kelleher and Markow 2007; Markow et al. 2007; Bono et al. 2011, 2015; Hardy et al. 2011; Richmond et al. 2012; Richmond 2014; McGirr et al. 2017; Diaz et al. 2021b; 2022)

The relatively close phylogenetic relationship to *D. melanogaster* has provided the *D. mojavensis* cluster with several advantages as a burgeoning genomic model system. The ability to detect chromosomal inversions via the analysis of polytene chromosomes, pioneered in *D. melanogaster*, allowed for early investigations into clinal variation as well as interpopulation and interspecific variation in inversions (Mettler 1963; Johnson 1980). More recently, the *D. mojavensis* population from Santa Catalina Island was among the first non-model *Drosophila* genomes sequenced (*Drosophila* 12 Genomes Consortium 2008), giving the species of the *D. mojavensis* cluster a high-quality starting point and a template for further research. Additionally, the wealth of functional genomic knowledge in *D. melanogaster* has allowed for clear interpretation of gene-level results as compared to more distantly related insects. This has been leveraged in a slew of candidate gene studies (Krebs 1999; Matzkin and Eanes 2003; Matzkin 2004; 2005; 2008; Guillén and Ruiz 2012; Diaz et al. 2018), whole-genome studies of molecular evolution (Guillén et al. 2015; 2019; Allan and Matzkin 2019; Rane et al. 2019), transcriptomics (Matzkin et al. 2006; Bono et al. 2011; Matzkin and Markow 2009; 2013; Matzkin 2012; Rajpurohit et al. 2013; Smith et al. 2013; Etges et al. 2015, 2017; Crowley-Gall et al. 2016; Nazario-Yepiz et al. 2017; Mateus et al. 2019; Benowitz et al. 2020; Banho et al. 2021ab; Diaz et al. 2021b; 2022), and functional analysis via CRISPR derived transgenics (Khallaf et al. 2020).

Despite this extensive history of genomic research, the data needed to address many key hypotheses within this system remains unavailable. At present, there is only a *de novo* sequenced genome for one of the four *D. mojavensis* populations, in spite of the outsized role that these populations have played in understanding molecular adaptation to variable host environments. Although this genome assembly is excellent, and has capably facilitated genetic mapping studies (Etges et al. 2007, 2009, 2010; Benowitz et al. 2019), several chromosomes, notably the X chromosome, remain far from contiguous. Outside of *D. mojavensis*, there are currently only two highly fragmented genome assemblies, one each from *D. navojoa* and *D. arizonae* (Sanchez-Flores et al. 2016; Vanderlinde et al. 2019).

Here, we take a major step towards addressing this gap and lack of genomic resources by first re-scaffolding and polishing the existing *D. mojavensis* genome from Santa Catalina Island, substantially improving this assembly. We then build *de novo* genome assemblies for strains from the other three *D. mojavensis* populations, two strains of *D. arizonae* collected from opposite ends of its range, and a single strain of *D. navojoa* (Fig. 1; Table 1). By using a hybrid assembly approach combining short- and long-read sequencing technologies, we are able to construct entire chromosomes for each genome, leading to some of the most complete assemblies throughout the *Drosophila* clade. We first use these assemblies to resolve longstanding questions regarding the phylogeny and divergence times within this group. We then assess protein-coding and structural evolution across all seven genomes. This allows us novel insight into the rates of each type of evolutionary divergence in this clade, and also provides the power to test fundamental hypotheses on the relationship between structural and coding evolution. Lastly, in order to facilitate the use of these genomes as a resource for the communities of *Drosophila* biologists and ecological geneticists, we present a public database of the assemblies and annotations (cactusflybase.arizona.edu).

**Table 1:** Information on stocks, from the National *Drosophila* Species Stock Center (Cornell) used for genome sequencing in this study.

| Species | Population Abbreviation | Location of Collection | Date of Collection | Stock Center ID | Local ID |
| --- | --- | --- | --- | --- | --- |
| *D. mojavensis* | BC | La Paz, Baja California Mexico | 2001 | 15081-1354.01 | MJBC 155 |
|  | CI | Santa Catalina Island, California US | 2002 | 15081-1352.22 | 15081-1352.22 |
|  | MOV | Anza-Borrego State Desert Park, California US | 2002 | 15081-1353.01 | MJANZA 402-8 |
|  | SON | Guaymas, Sonora Mexico | 1998 | 15081-1355.01 | MJ 122 |
| *D. arizonae* | ARI | Guaymas, Sonora Mexico | 2004 | 15081-1271.41 | AR002 |
|  | CHI | Chiapas, Mexico | 1987 | 15081-1271.14 | AZ Chiapas 1B 13610 |
| *D. navojoa* | NAV | Jalisco, Mexico | 1997 | 15081-1374.11 | 15081-1374.11 |

**Figure 1.**
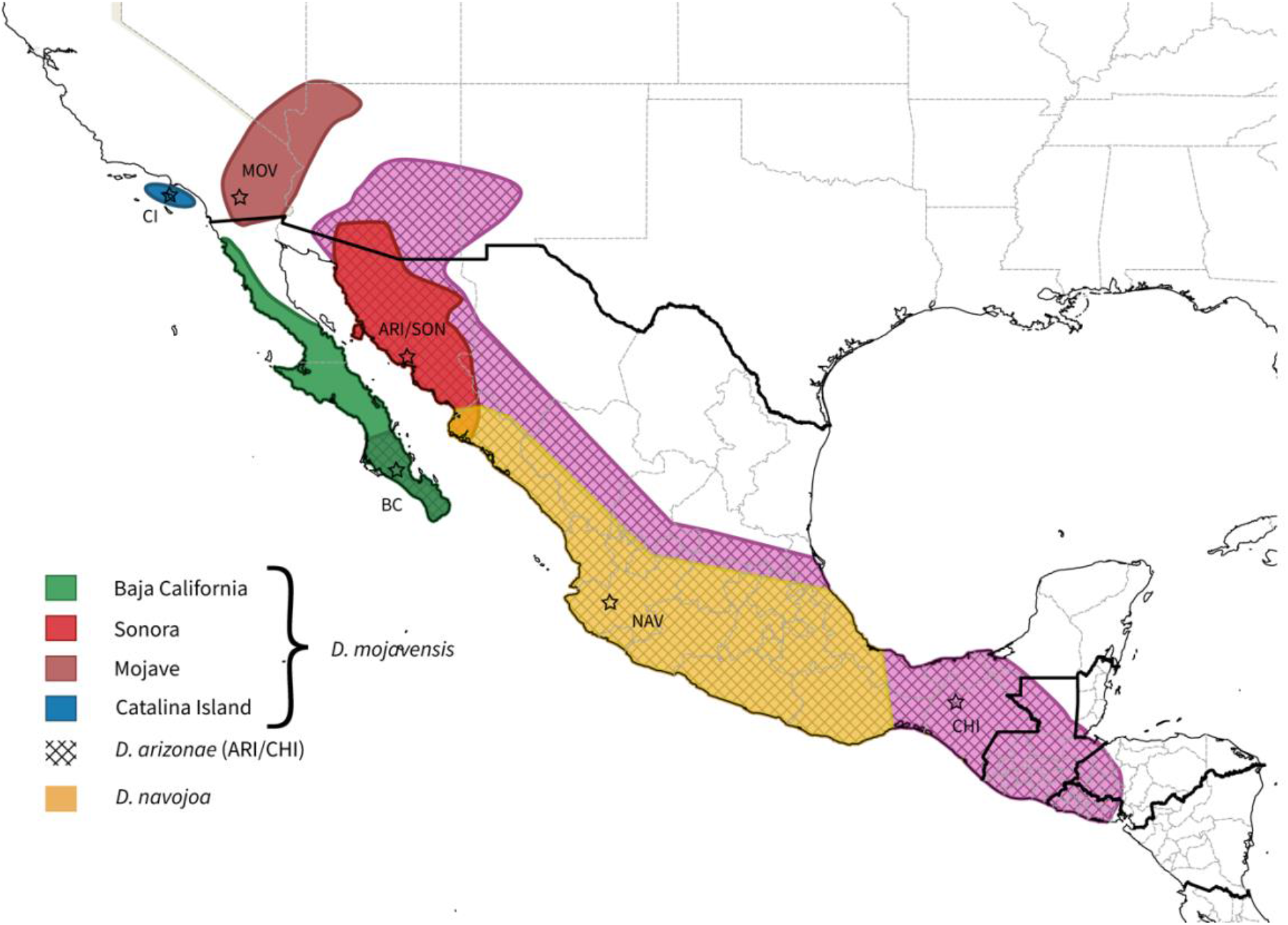
Ranges of the species and populations sequenced in this study. Hatched regions represent the range of *D. arizonae*. No discrete geographical boundary is known to separate the ARI and CHI populations sequenced here. Stars show the location of collection of the genome lines. Ranges are estimated based on collection site and host plant ranges.

## Results

### Genome assembly and annotation

Details of the strains used for genome sequencing can be found in Table 1. Genome assembly and annotation statistics can be found in Table 2. All six *de novo* assemblies were highly contiguous, with all six major chromosomes assembled with only a handful of gaps. The re-scaffolding of the original CI genome also resulted in higher contiguity, although the assembly still has a higher percentage of gaps as well as repeats, which could indicate the presence of redundant scaffolds.

**Table 2:** Genome assembly statistics

|  | CI | MOV | BC | SON | ARI | CHI | NAV |
| --- | --- | --- | --- | --- | --- | --- | --- |
| Genome size (Mb) | 191.84 | 160.64 | 161.282 | 158.92 | 163.52 | 162.67 | 156.70 |
| # of scaffolds | 6,327 | 69 | 68 | 42 | 45 | 39 | 68 |
| Scaffold N50 length (Mb) | 32.37 | 32.40 | 32.47 | 32.28 | 33.82 | 33.67 | 31.27 |
| # of contigs | 10,611 | 71 | 69 | 46 | 51 | 43 | 69 |
| Contig N50 length (Mb) | 0.041 | 27.01 | 27.38 | 26.92 | 27.42 | 27.17 | 27.05 |
| Gaps (Mb) | 12.26 | 0 | 0 | 0 | 0 | 0 | 0 |
| GC content (%) | 39.48 | 39.66 | 39.67 | 39.64 | 39.7 | 39.65 | 39.95 |
| Repeat content (%) | 29.27 | 25.19 | 25.48 | 24.5 | 26.48 | 26.23 | 23.79 |
| Number of proteins | 13,755 | 13,358 | 13,388 | 13,426 | 13,408 | 13,321 | 13,203 |
| Genome BUSCO (%) |  |  |  |  |  |  |  |
| Complete | 99.0 | 99.1 | 99.0 | 99.1 | 99.1 | 99.1 | 99.2 |
| Single copy | 98.6 | 98.8 | 98.6 | 98.8 | 98.8 | 98.8 | 98.8 |
| Duplicated | 0.4 | 0.3 | 0.4 | 0.3 | 0.3 | 0.3 | 0.4 |
| Fragmented | 0.4 | 0.3 | 0.5 | 0.4 | 0.2 | 0.3 | 0.4 |
| Missing | 0.6 | 0.6 | 0.5 | 0.5 | 0.7 | 0.6 | 0.4 |
| Proteome BUSCO (%) |  |  |  |  |  |  |  |
| Complete | 99.3 | 98.9 | 98.8 | 99.0 | 98.9 | 98.9 | 99.1 |
| Single Copy | 98.8 | 98.4 | 98.4 | 98.5 | 98.4 | 98.4 | 98.6 |
| Duplicated | 0.5 | 0.5 | 0.4 | 0.5 | 0.5 | 0.5 | 0.5 |
| Fragmented | 0.3 | 0.5 | 0.5 | 0.5 | 0.5 | 0.5 | 0.4 |
| Missing | 0.4 | 0.6 | 0.7 | 0.5 | 0.6 | 0.6 | 0.5 |

Genome size was consistent across the three new *D. mojavensis* genomes, with both *D. arizonae* exhibiting larger genomes and *D. navojoa* slightly smaller. There is no evidence that these evolutionary patterns in genome size are driven by expansions of repeats or TEs, as these did not display such consistent trends between the species.

Our genome assemblies confirmed previous findings on fixed chromosomal inversions between these species and populations (Fig. 2; Supplemental Figs. S1, S2), with a single inversion occurring at the base of *D. mojavensis* on Muller element A (X chromosome), and multiple overlapping inversions on Muller elements B and E. Some inversion breakpoints were associated with windows of high repeat and TE content (Fig. 3; Supplemental Fig. S3), but this pattern was not ubiquitous.

**Figure 2.**
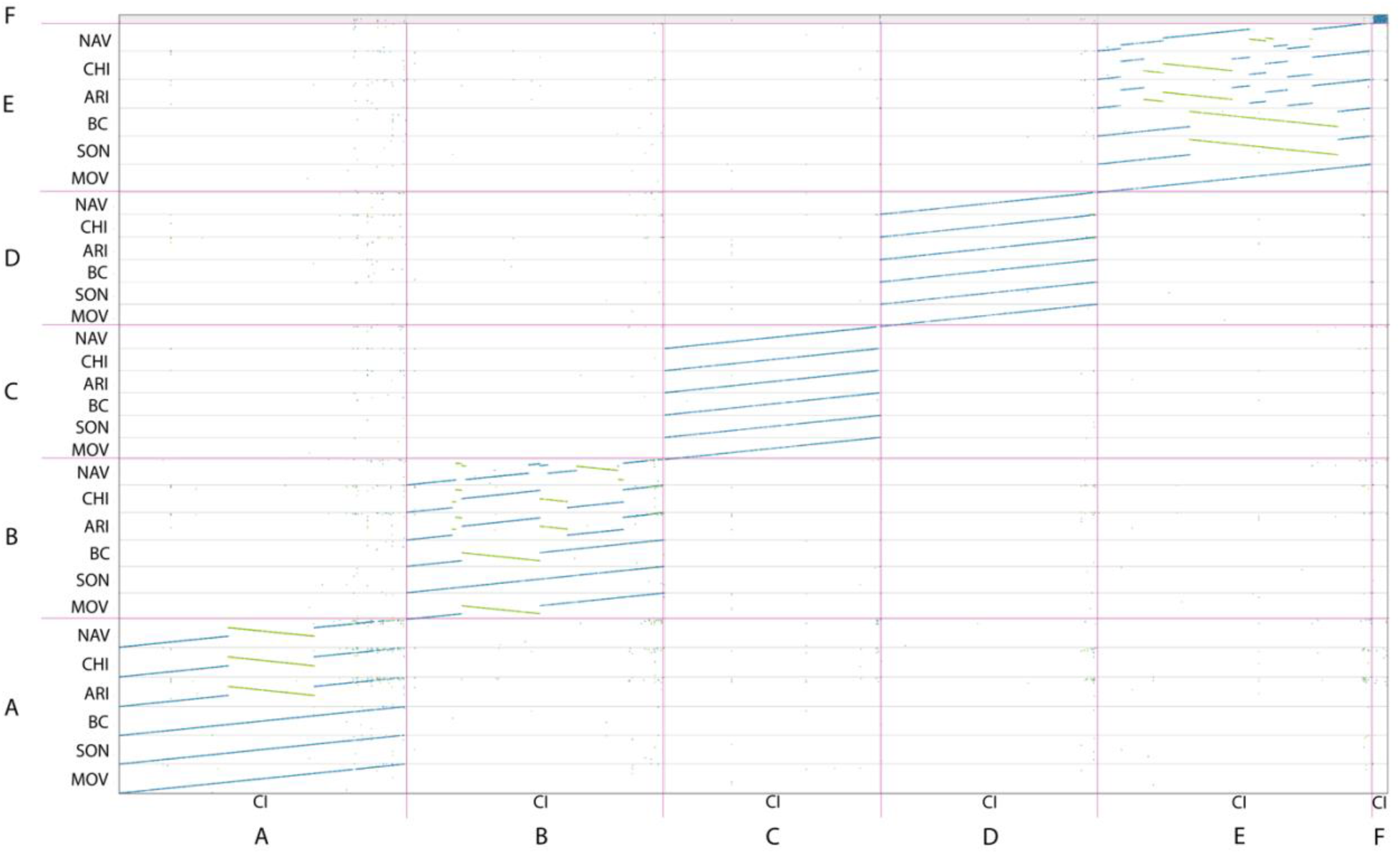
Syntenic conservation of the six *de novo* genomes sequenced in this study as compared to the reassembled CI genome. Letters A-F indicate Muller elements. Green lines indicate inversions.

**Figure 3.**
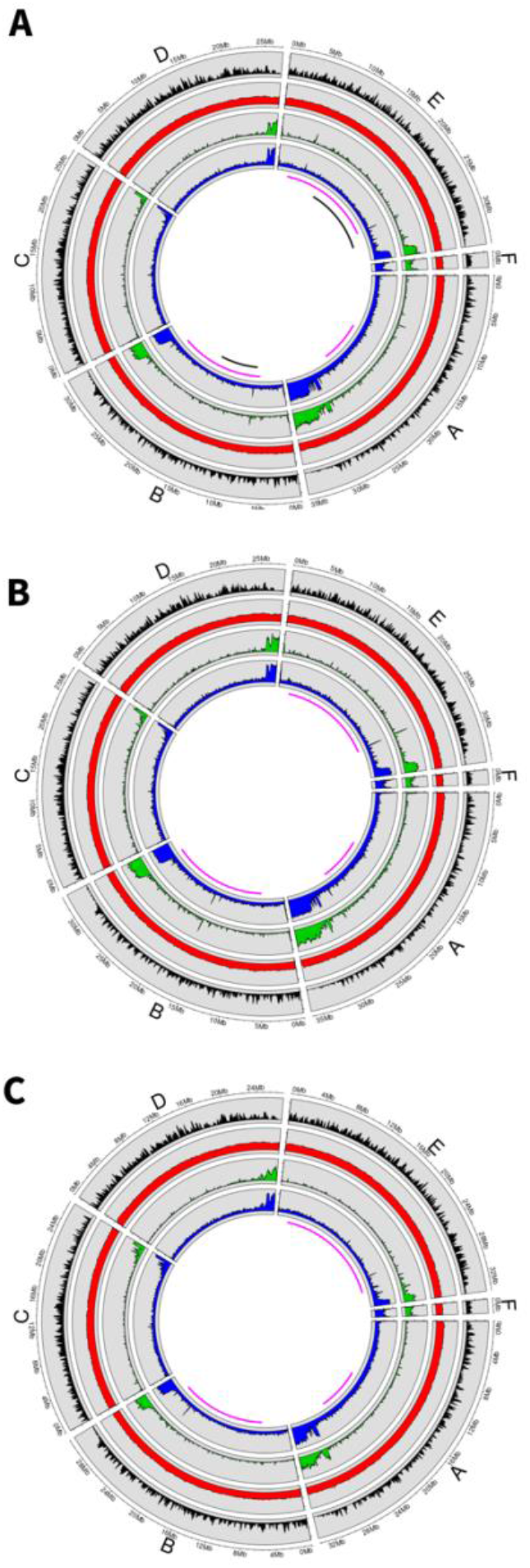
Genome statistics for: (*A*) Mojave *D. mojavensis*, (*B*) Chiapas *D. arizonae*, and (*C*) *D. navojoa*. From outside to in, circles represent gene content, GC content, TE content, and total repeat content. Pink bars below the circles represent the regions covered by interspecific inversion polymorphisms, and black bars represent regions covered by inversion polymorphisms within *D. mojavensis*.

To facilitate further study of these species, we have deposited the assemblies and annotations in a new public database at cactusflybase.arizona.edu. Users can download fasta and gff files directly, view annotations and underlying RNA-seq data via JBrowse (Buels et al. 2016), and BLAST the genome and proteome databases using SequenceServer (Priyam et al. 2019). Details of the species and populations sequenced and their husbandry are available as well.

### Phylogenomics and divergence time estimation

Both the topology of the phylogeny as well as the divergence time estimates differed when using nuclear (Fig. 4) versus mitochondrial genes (Supplemental Fig. S4). The nuclear derived phylogeny placed the four *D. mojavensis* populations in a single clade, with the two *D. arizonae* populations as a sibling clade, and *D. navojoa* as an outgroup. While the mitochondrial phylogeny also had *D. navojoa* as an outgroup, it included the northern *D. arizonae* population as part of the *D. mojavensis* clade, with the Chiapas population an outgroup to that clade. Both phylogenies agreed that the BC and SON *D. mojavensis* populations were most closely related, although other aspects of the topology within *D. mojavensis* also differed.

**Figure 4.**
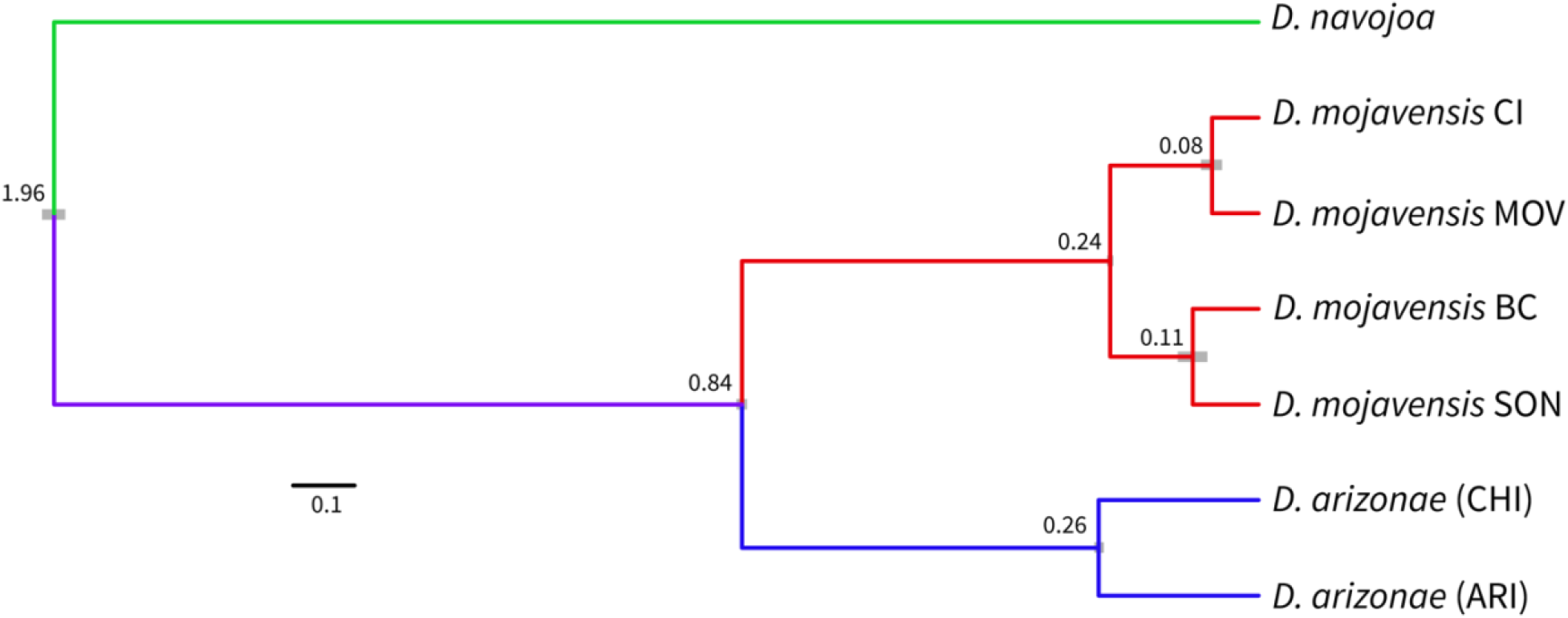
Phylogeny and divergence times (mya) as estimated by 12,218 single copy nuclear genes. Colors represent the accepted species identities and grey bars represent 95% confidence intervals for divergence time estimates.

Although the timing of divergence within the *D. arizonae*/*D. mojavensis* clade was identical between the two datasets, the mitochondrial phylogeny gave a twice as old split of *D. navojoa* from this group.

### Syntenic evolution

We defined syntenic (or collinear) regions of the genome as those displaying one-to-one conservation of sequence as called by SyRI (Goel et al. 2019) and syntenic divergence as the percentage of non-syntenic genome content between two genomes in a given region. Syntenic divergence of each of the six *de novo* sequenced populations from CI recapitulated the divergence patterns as found in the nuclear phylogeny. As expected, *D. navojoa* had by far the greatest mean syntenic divergence, while both *D. arizonae* populations had nearly identical levels of divergence. MOV had slightly higher synteny compared to BC and SON. Overall, the breakdown of synteny over evolutionary time was found to be linear, with a loss of roughly 33.67% of genome collinearity per million years (Supplemental Fig. S5)

Independently of chromosomal inversions, which were rearranged in all seven genomes to match the CI karyotype prior to analysis, significant variation in syntenic divergence was present between chromosomes. Muller elements A and F showed reduced synteny in all six genomes. In *D. arizonae*, Muller elements B and E, which also carry inversions, displayed lower synteny than C and D, which do not carry inversions. Patterns in *D. navojoa* were similar apart from a reduction in synteny on Muller element C compared to D (Fig. 5A-G).

**Figure 5.**
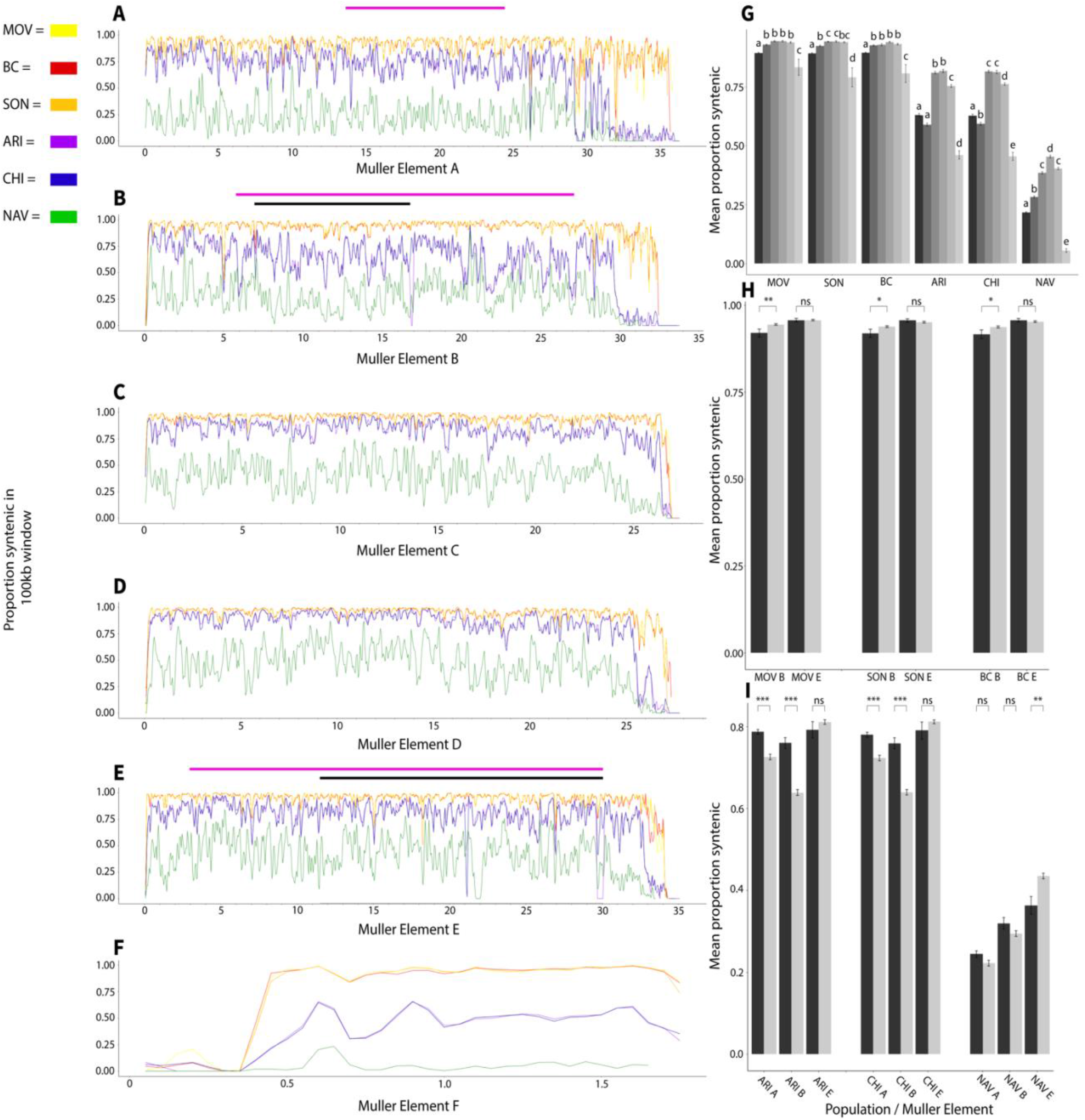
(*A-F*) The synteny score across 100kb windows for Muller elements A-F. MOV, yellow; SON, orange; BC, red; ARI, purple; CHI, blue; NAV, green. Pink and blue bars above plots indicate inter- and intraspecific inversion polymorphisms as in Figure 2. Numbers on x axis indicate position in Mb. (*G*) Mean synteny score for each element and species. Letters indicate significant differences (p < 0.05) between elements for each species. Muller elements are arranged in order with A (darkest) at left and F (lightest) at right. (*H*) Mean synteny scores before (dark grey) and within (light grey) inverted regions of elements B and E for the three *D. mojavensis* populations. (*I*) Mean synteny scores before (dark grey) and within (light grey) inverted regions of elements A, B, and E for the *D. arizonae* and *D. navojoa* genomes. Asterisks in parts (*H*) and (*I*) indicate significance at the level of p < 0.05 (*), p < 0.001 (**) or p < 0.0001 (***).

Within chromosomes bearing inversions, the relative rates of syntenic divergence inside and outside (measured here as only the region on the centromeric side of the inversion due to low synteny near telomeres) the inversion depended on the evolutionary distance and specific chromosome. Within *D. mojavensis*, Muller element B displayed greater divergence prior to the chromosomal inversion breakpoint (we did not compare regions after the inversions due to major reductions of synteny in telomeres; Fig. 5A-F) in all three populations, including the SON population, which is homokaryotypic with CI (Fig. 5H). However, synteny in Muller element E was consistent before and within the inversion. Interspecific syntenic divergence, on the other hand, was greater within the inversion regions on Muller elements A and B, while the opposite was true on E (Fig. 5I).

### Molecular evolution

Lists of genes found to be under positive selection via BUSTED and codeml analyses can be found in Supplemental Tables S1 and S2. A comparison of gene families previously hypothesized to be involved in adaptation to variable cactus environments showed no elevated rates of positive selection in these gene families via the codeml analysis (Fig. 6). However, higher rates of positive selection were found within reproductive genes as well as orphan genes absent from *D. melanogaster*.

**Figure 6.**
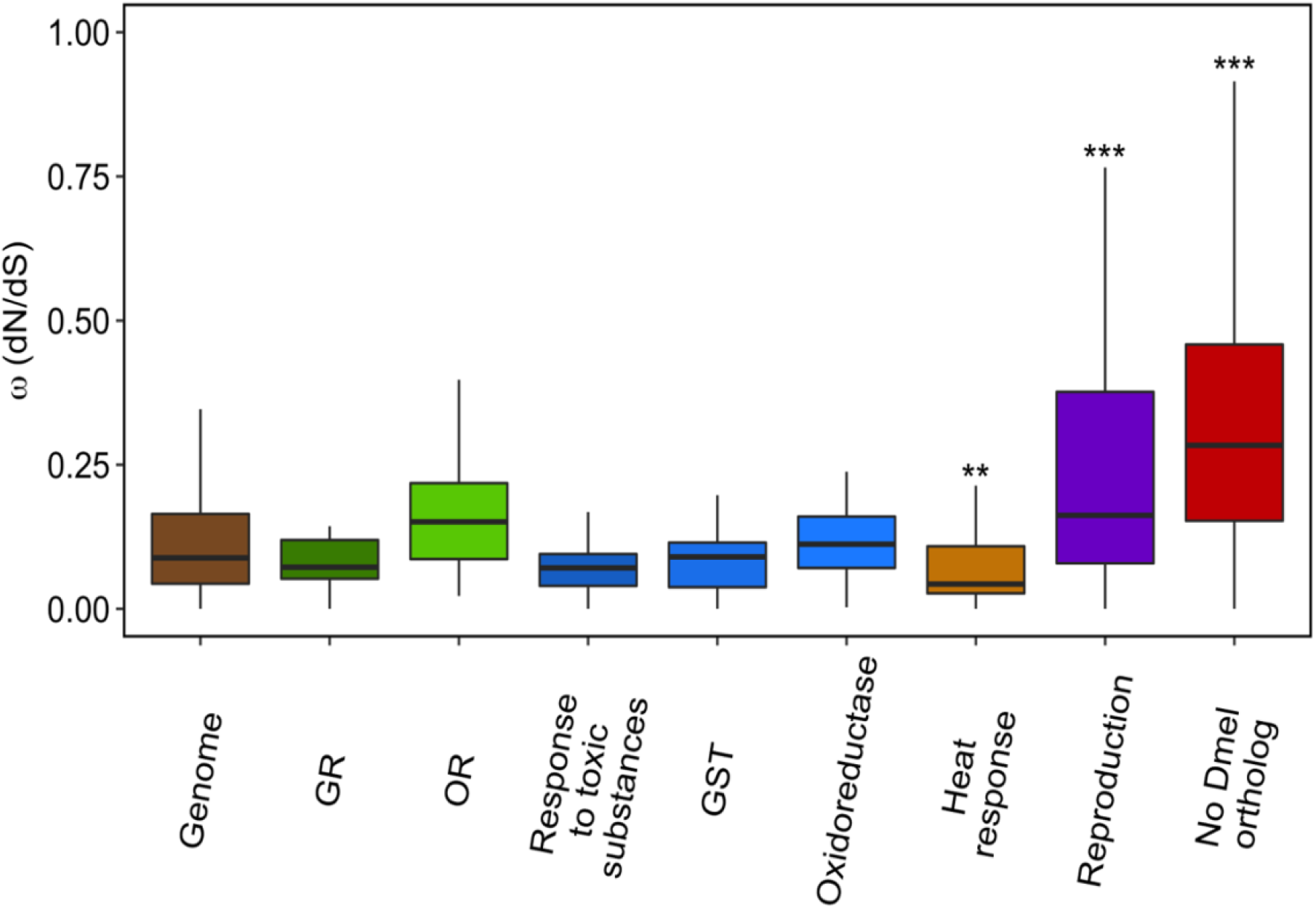
Comparison of omega values for different gene families and GO categories. Asterisks indicates significance at the level of p < 0.001 (**) or p < 0.0001 (***).

We found no evidence that genes surrounding either the breakpoints within *D. mojavensis* (F_1,12185_ = 0.017, p = 0.68) nor the breakpoints in the clade as a whole (F_1,12185_ = 0.33, p = 0.57) displayed elevated evolutionary rates. Rates of positive selection were significantly negatively corelated to the synteny score from CI to NAV of the sliding window containing the gene (Fig. 7). This pattern held for the synteny from CI to the mean of the *D. arizonae* populations (F_1,12185_ = 44.80, p = 2.28 × 10^−11^) as well as the synteny from CI to the mean of the other three *D. mojavensis* populations (F_1,12185_ = 19.89, p = 8.26 × 10^−6^).

**Figure 7.**
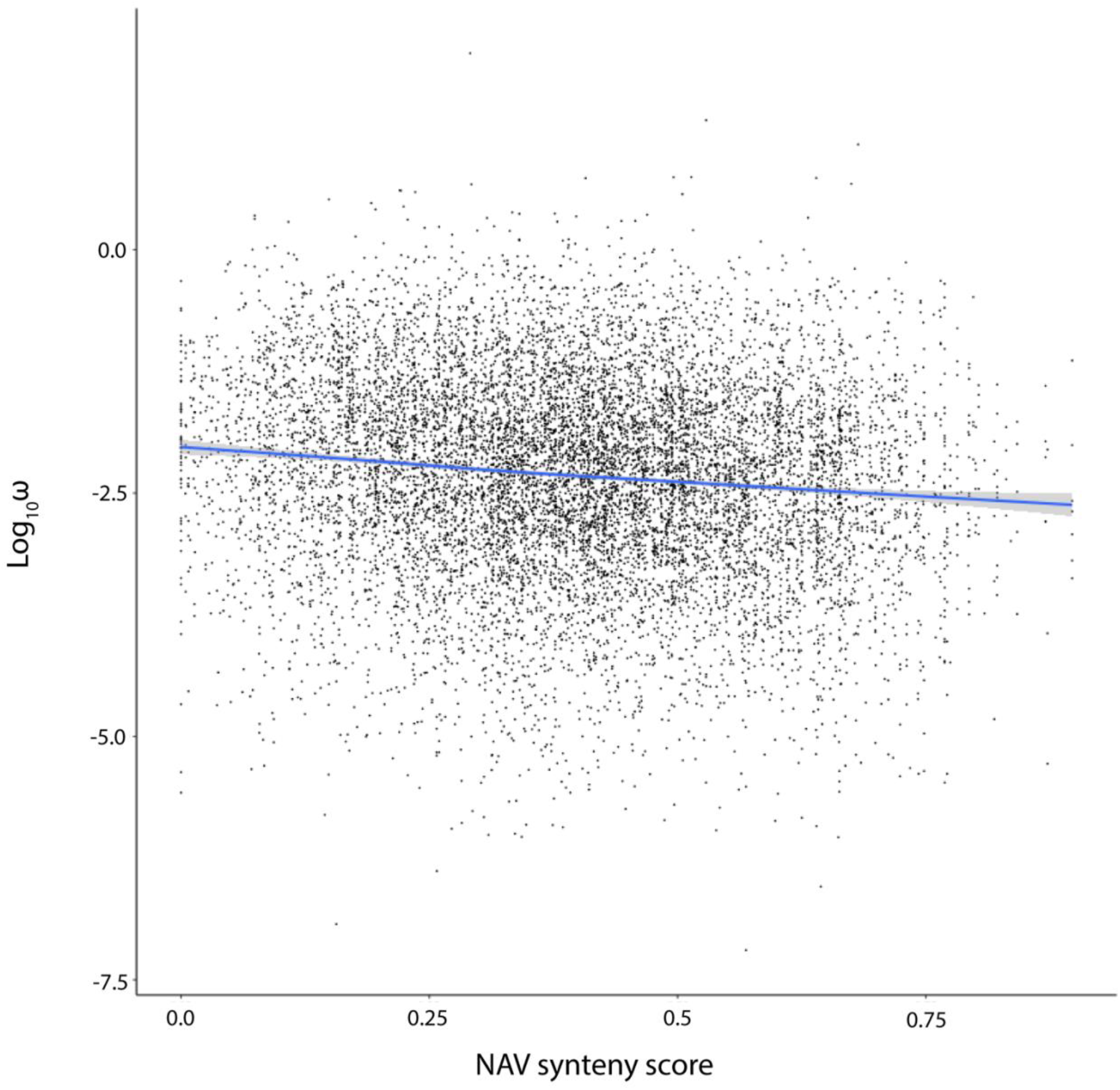
The genome-wide relationship between synteny (from *D. mojavensis* Catalina Island to *D. navojoa*) and molecular evolutionary rate across the phylogeny. The regression line represents a linear regression (y = -0.36x – 0.98), with 95% confidence intervals shaded (F_1,12153_ = 94.64, p < 2.2×10^-16^).

## Discussion

The assembly of these seven *de novo* genomes represents one of the largest and most contiguous genomic datasets in any *Drosophila* clade. To date, we can identify 21 genomes that have been assembled at or near chromosome level in *Drosophila* (https://www.ncbi.nlm.nih.gov/genome/browse#!/overview/Drosophila). However, these are heavily clustered in clades closely related to *D. melanogaster* and *D. pseudoobscura*. While there has been a clear logic to concentrating sequencing effort on a small handful of clades, the establishment of generalizable patterns of genomic evolution requires that analyses be replicated across taxa. Our genomes add to this endeavor. Furthermore, the inclusion of separate conspecific populations of two species allows for deeper resolution in the calculation of fundamental evolutionary patterns.

Accurate phylogenies and divergence times are critical both for quantifying rates of evolutionary change and for generating hypotheses on the phylogeographic causes of speciation events and adaptive radiations. Despite the extensive molecular investigation of the *D. mojavensis* species cluster, disagreement on both the topology and node ages of the phylogeny persists. Within *D. mojavensis*, three different trees have been supported. A mitochondrial study (Reed et al. 2007) found the Mojave Desert population as an outgroup while a nuclear study (Smith et al. 2012) placed the Baja California population as an outgroup. Two earlier nuclear studies (Ross and Markow 2006; Machado et al. 2007) found two clades, with Mojave - Catalina Island and Baja – Sonora as pairs of sibling species. Our mitochondrial data recapitulated the topology of the earlier mitochondrial tree, while our nuclear data supported the topology of the earlier studies (Ross and Markow 2006; Machado et al. 2007). These studies also presented variable divergence times; nuclear studies (Machado et al. 2007; Smith et al. 2012) found the initial divergence within *D. mojavensis* to have occurred ∼250,000 years ago with further divergence between 100,000 and 150,000 years ago, the mitochondrial study found node ages more than twice as old. Once again, our mitochondrial and nuclear data cleanly recover these differences.

Our mitochondrial and nuclear analyses also showed major differences in the relationships and divergence times across species. The biggest among these is the finding of paraphyly in *D. arizonae* in our mitochondrial dataset. Although one analysis in previous mitochondrial work (Reed et al. 2007) also failed to support *D. arizonae* as a clade, it differed in which population grouped with *D. mojavensis*. Here, our patterns of syntenic divergence align with the nuclear dataset in grouping the two *D. arizonae* populations as sibling taxa.

These consistent differences between mitochondrial and nuclear datasets could reflect a different demographic history for the mitochondrial genome, as previously suggested (Reed et al. 2007). However, we consider this discrepancy to be more likely due to noise, given the nearly 10,000-fold difference in genes analyzed (12,218 vs. 13) in the nuclear genome compared to the mitochondrial genome. Thus, our data best support the topology of the earlier nuclear phylogeny of Machado et al. (2007), with divergence times broadly in agreement with both previous nuclear studies (Machado et al. 2007; Smith et al. 2012).

Our results regarding divergence times between species significantly contradicted estimates from earlier work. These estimates have ranged widely, ranging from 0.66 mya to 4.2 mya for the split between *D. mojavensis* and *D. arizonae*, and from 2.9 to 7.8 mya for the divergence of these species to *D. navojoa* (reviewed in Sanchez-Flores et al. 2016). In the most robust analysis to date, Sanchez-Flores et al. (2016) used over 5,000 nuclear loci to estimate an age of 5.86 mya for the split between *D. navojoa* and the rest of the *D. mojavensis* cluster and an age of 1.51 mya for the split between *D. arizonae* and *D. mojavensis*. Our nuclear analysis showed much younger divergence times of 1.96 mya for the split of *D. navojoa* and 0.84 mya for the split of *D. arizonae*. We argue that our results are more reliable for three reasons. First, our analysis utilized more than twice as many loci, as we analyzed more than 12,000 single copy orthologs. Second, our usage of multiple genomes for both *D. mojavensis* and *D. arizonae* further reduced the possibility for sampling error based on analyzing single genotypes per species. Third, our usage of the neutral mutation rate to calibrate the phylogeny is expected to be more accurate than using models calibrated from Hawaiian *Drosophila*, which have been found to inflate divergence times dramatically (Obbard et al. 2012). These younger estimates suggest that the speciation of this entire clade revolves around the cyclic climatic fluctuations of the past few million years and the accompanying shifts in host cactus distribution. On the contrary, major geological events such as the raising of the Trans-Mexican Volcanic Belt, that have been previously hypothesized as possible causes of intra- and interspecific divergence (Machado et al. 2007; Rampasso et al. 2017), appear to be too ancient to have played a role here.

Descriptions of the rates of sequence and expression evolution have served as foundational patterns of evolutionary genomics for decades. However, limited data relating to rates of accumulation of structural genomic variation have been published. Chakraborty et al. (2021) found that 15% of sequence did not align between *D. simulans* and *D. melanogaster*, which are diverged by about 3 million years, and noted that this was over twice the percentage of sequence variation between these species. Long et al. (2018) estimate a rate of 50 structural mutations per Mb per million years within *D. melanogaster*, which, given an average variant size of around 25 kb in their dataset, amounts to approximately 13% divergence per million years. Jiao and Schneeberger (2020) report around 10% syntenic divergence between *Arabidopsis thaliana* accessions using the same software and methodology here, but cannot present a phylogenetic timeline of breakdown. Here, we observe that synteny decays in a linear fashion, with about 33% of genome collinearity lost per million years. Although this conclusion is undoubtedly sensitive to the particular parameters used to define a syntenic block, we hope this can serve as a baseline for future studies quantifying the evolution of genome structure both within *Drosophila* as well as in other taxa. Given increasing implication of structural variants in speciation (Zhang et al. 2021) and adaptation (Mérot et al. 2020), we are curious to see how these results stand out in an even broader comparative framework. Is this a fast or slow rate of structural divergence? Do structural variants accumulate more rapidly in highly speciose taxa, or those undergoing adaptive radiations?

To begin to address these questions within our dataset, we asked what factors might explain local variation in synteny within the *D. mojavensis* group. The best predictor of syntenic divergence was chromosome. Although results varied slightly depending on the evolutionary distance, the dot chromosome (Muller element F) diverged most rapidly, followed by Muller elements A, B, and E. In nearly all comparisons, Muller elements C and D maintained the greatest collinearity. It is unlikely that this heterogeneity can be explained by a single factor. For Muller element F, although there is some evidence for relaxed constraint in *D. mojavensis* (Allan and Matzkin 2019), it is more likely that our results are explained by a genus-wide propensity for this chromosome to accumulate repeats and TEs, which has been attributed to a unique chromatin structure for this chromosome (Riddle and Elgin 2018). The consistent degradation of the X chromosome, on the other hand, appears to be linked to increased repeat but not TE content. This breakdown may be linked to the prevalence of rapidly evolving tandem repeats known to be common on *Drosophila* X chromosomes (Sproul et al. 2020).

No such variation in TE or repeat content is apparent amongst the four large autosomes. Instead, the variation in collinearity of these chromosomes is noteworthy for its association with the presence of major inversions. Muller elements B and E have inverted repeatedly in the *D. mojavensis* cluster, including multiple times at nearly identical breakpoints, whereas C and D have not (Supplemental Fig. S2). Both adaptive and neutral hypotheses have been considered for the reuse of breakpoints. Adaptive explanations have focused on the potential for inversions to prevent recombination across genes involved in local adaptation, therefore maintaining positive combinations of alleles together (Hoffman et al. 2004; Kirkpatrick and Barton 2006; Wellenreuther and Bernatchez 2018). Non-adaptive explanations have considered that certain genomic regions may be susceptible to inversions due to variation in chromatin structure and genome fragility (von Grotthuss et al. 2010). Our results support the latter explanation for the *D. mojavensis* cluster, as Muller elements B and E appear to be more susceptible to a wide range of structural mutations beyond large inversions. Further supporting that this relationship is correlational, we see no evidence that inversions cause additional decreases in synteny, as there was no consistent trend of increased collinearity outside of the inverted regions of these chromosomes. This does not exclude the possibility that specific breakpoints are relevant to adaptation; although we found no evidence that genes near breakpoints within the *D. mojavensis* cluster are more likely to display signatures of selection, the presence of some positively selected genes near breakpoints still reflects a potential link between inversions and adaptation. Furthermore, previous work (Guillén and Ruiz 2012) suggests that gene regulatory variation may be responsible for inversion associated adaptation in this system.

We also found that variation in synteny was negatively linked to omega. In other words, genes with greater rates of protein-coding evolution were more likely to occur in regions of decreased collinearity. Two nonexclusive phenomena could help explain this pattern. First, genes already experiencing relaxed selection on protein function might better tolerate structural changes that may also influence splicing or expression changes (Hämälä et al. 2021), meaning that mutations near these genes are more likely to be maintained. Second, the causality could be reversed, and structural changes to genes could directly cause subsequent bouts of reduced constraint and relaxed selection. In many cases, this could be explained as a result of sub- or neofunctionalization following gene duplication. However, in our dataset, molecular evolution was only assessed for single copy orthologs across all seven genomes. Thus, the relevant duplications would have occurred prior to the common ancestor of these species, and would not register as structural variants in this dataset. A more likely possibility is that structural changes resulting in alterations to gene regulation alter protein function, which subsequently leads to a relaxation of purifying selection on amino acid sequences.

Given this relationship, we consider genes with elevated rates of positive selection in regions of low collinearity to be especially strong candidates for roles in adaptation and speciation. We are particularly interested in genes involved in reproduction, given the elevated rates of positive selection for genes in this category. One particularly interesting gene in this regard is GI18186, which has an omega of 1.166 and lies in a window with a synteny score in the 6^th^ percentile or lower in all three species. This gene is orthologous to the *D. melanogaster* gene CG13965, which is massively expressed in male accessory glands (Brown et al. 2014) and has been localized to a small cluster of accessory gland proteins (Acps; Ravi Ram and Wolfner 2007). Furthermore, CG13965 protein is known to be transferred from males to females during mating, not only in *D. melanogaster* (Immarigeon et al. 2021) but in *D. simulans* and *D. yakuba* as well (Findlay et al. 2008). Function of male protein in the female reproductive tract has been hypothesized as an important speciation mechanism between species and populations in the *D. mojavensis* cluster (Bono et al. 2011). Our results suggest that GI18186 is worthy of further attention, and that both changes to the expression and sequence of this gene may have contributed to pre-mating post-zygotic isolation leading to reproductive isolation, as is the case for Acps in *D. melanogaster* (Immarigeon et al. 2021). Given that the number of annotated Acps in *Drosophila* is in the hundreds, it is important to narrow down the list of possible relevant genes for more targeted studies. Thus, it is valuable that our integration of sequence and structural analysis allows us to make this prediction from single genome sequences alone.

Extending this, the second category of genes that were found to be overrepresented for positive selection are those without orthologs in *D. melanogaster*, and are therefore likely to be taxonomically restricted genes (TRGs) in at least the *repleta* group if not the *D. mojavensis* cluster. TRGs have been previously implicated in cactophilic *Drosophila* evolution (Moreyra et al. 2022) as well as many other taxa, and likely reflects both that TRGs are unlikely to have housekeeping functions and may be preferentially involved in novel traits and adaptations (Domazet-Loso and Tautz 2003; Arendsee et al. 2014; Jasper et al. 2015). In spite of their likelihood of relevance to adaptation, the lack of functional annotation for genes with no well-studied ortholog in a model organism represents a major issue in the biology of non-model organisms, and a systematic study of these genes is unlikely for the vast majority of taxa. Here, we find that most of the genes with evidence of positive selection in regions of low collinearity are TRGs. We argue that these genes should be prioritized in targeted investigations seeking to characterize the functions of currently unstudied genes.

## Materials and Methods

### Insect strains, genome sequencing, and assembly

Each strain used in this study (Table 1) was maintained as an inbred line in the Matzkin lab at the University of Arizona on a banana-molasses based diet (recipe in Coleman et al. 2018) through genome and RNA sequencing.

The original genomic scaffolds (*Drosophila* 12 Genomes Consortium 2007) as well as short- and long-read (Miller et al. 2018) sequence data for the Santa Catalina Island *D. mojavensis* assembly are previously published. The short-read Illumina data for the remaining *D. mojavensis* populations is described in Allan and Matzkin (2019). Long-read data for the Sonora *D. mojavensis* population is described in Jaworski et al. (2020). The short-read Illumina data for the *D. arizonae* Sonora population is described in Diaz et al. (2021b). The short-read Illumina data for *D. navojoa* is described in Vanderlinde et al. (2019).

Briefly, for all short-read data, we extracted DNA from a pool of ten adult males and ten adult females using Qiagen DNeasy Blood & Tissue Kits (Qiagen, Hilden, Germany), and we constructed the *D. arizonae* Chiapas library using KAPA LTP Library Preparation Kit (Roche, Basel, Switzerland) kits. It was sequenced on an Illumina HiSeq 4000 at Novogene (Beijing, China) at 220X coverage. All other short-read libraries were built and sequenced on Illumina HiSeq 2000 at the HudsonAlpha Genome Sequencing Center (Huntsville, AL, USA) at 75X coverage. For all long-read data, we extracted high molecular weight DNA from a pool of 150 males and 150 females using a chloroform-based extraction, detailed method in Jaworski et al. (2020). PacBio libraries were built and sequenced on a PacBio Sequel at the Arizona Genomics Institute (Tucson, AZ, USA).

The assembly of the six *de novo* genomes largely followed the hybrid assembly strategy described in Jaworski et al. (2020), wherein a detailed description of sequencing and assembly methods can be found. Briefly, we used Platanus 1.2.4 (Kajitani et al. 2014) and DBG2OLC (Ye et al. 2016) to produce hybrid assemblies of the short- and long-read data. We also used Canu 1.7 (Koren et al. 2017) for long-read only assembly with the correctedErrorRate parameter set to 0.039 for the primary assembly though this was increased to 0.065 to produce a less stringent assembly used for bridging and extending primary contigs. We used Quickmerge 2.0 (Chakraborty et al. 2016) to merge these two assemblies into a draft assembly. We then manually merged contigs based on whole genome alignments from Mauve (Darling et al. 2004) and Nucmer (Delcher et al. 2002) including using the less stringent assembly in Geneious Prime (Biomatters, Auckland, NZ). Where contigs could not be merged, we manually joined them based on alignment with the other genomes and connected with an N-gap of 100bp. We aligned all remaining contigs not assigned to a chromosome with Minimap2 (Li 2018) and subsequently discarded all contigs with a match of over 80% to a chromosome scaffold. We polished each genome three times with Pilon 1.23 (Walker et al. 2014). During manual curation of our annotations with the help of RNA-seq data (see below), we identified several small insertion/deletion errors in each genome that led to frameshift errors causing problems with gene structure, and subsequently fixed these errors manually in Geneious Prime. We noticed that the *D. arizonae* Chiapas genome had substantially more of these errors than the others and therefore polished it a fourth time with Pilon 1.23 before fixing remaining errors manually as for the other genomes. We also performed additional polishing of the gene-containing regions of *D. navojoa* using majority consensus in Geneious Prime.

We reassembled the *D. mojavensis* Santa Catalina Island genome (hereafter, CI) in order to provide better comparisons with the six *de novo* assemblies. We first polished the most current FlyBase assembly (version r1.04) twice with Pilon 1.23. We then manually scaffolded by aligning contigs from the existing Nanopore data (Miller et al. 2018) to the polished reference using Mauve and joining in Geneious Prime. We filled all N-gaps over 20kb with contigs from the Nanopore dataset. Lastly, we filled all N-gaps regardless of size if they occurred within 100bp of a putative CDS feature identified during annotation. Similar to the other assemblies, annotation revealed several indel errors in coding regions the CI genome, which we fixed manually. In addition, to filtering duplicate scaffolds with Minimap2 we also removed scaffolds that previously had a gene annotation in the 1.04 release if those genes had strong BLAST hits to a gene on the chromosome scaffolds. Existing annotations were kept if no BLAST hit was found, all other annotations on unmapped scaffolds were removed.

We noticed that the previous assembly of Muller element F in CI was much larger than in our *de novo* assemblies, and contained ∼1.3Mb of sequence that was syntenic to sequence in Muller element A (X chromosome). We therefore split the CI Muller element F into two pieces: we kept bp 1-2,135,734 as chromosome F, while we joined bp 2,139,764-3,406,379 to chromosome A based on alignments in Mauve and NUCmer. We confirmed this split based on separate mapping data from a cross of the CI, SON, and MOV *D. mojavensis* populations, which showed no genetic linkage across this breakpoint of the original chromosome F (K.M. Benowitz unpubl. Data). All other large scaffolds in CI were linked to a chromosome based on physical and genetic marker data from Schaeffer et al. (2008).

After finalizing the assemblies, we ran RepeatModeler (Flynn et al. 2020) on each genome before using USEARCH (Edgar 2010) with a 90% similarity cutoff to create a non-duplicated combined list of repetitive elements. We then ran RepeatMasker (http://www.repeatmasker.org) to generate masked versions of each assembly prior to annotation.

We generated mitochondrial assemblies for all six *de novo* genomes by mapping reads to the existing CI mitochondrial sequence (*Drosophila* 12 Genomes Consortium 2008) in Geneious Prime.

### Genome annotation

To help facilitate annotation, we performed a broad RNA-seq experiment designed to detect expression of as many genes as possible. In October 2020, we collected tissue from each of the seven genome strains during early (12 hours post-laying) and late (26 hours post-laying) embryonic stages, first, second, and third instar larvae, pupae, and male and female adults at varying ages post-eclosion. For each life stage, we ground tissue in 500 μL of Trizol reagent (Thermo Fisher Scientific, Waltham, MA, USA) prior to extracting RNA using a ZYMO Direct-zol RNA Miniprep Kit. We then quantified the RNA and pooled extractions for each life stage together to reach 1.5 μg of total RNA. We then built libraries using a KAPA stranded mRNA-Seq Kit for each strain and sequenced them on an Illumina HiSeq 4000 lane at Novogene. We trimmed all RNA reads using Trimmomatic (Bolger et al. 2010) and aligned each to its respective genome using Hisat2 (Kim et al. 2019) under default parameters.

We used the current annotation of the Catalina Island *D. mojavensis* genome (*Drosophila* 12 Genomes Consortium 2008) as a starting point for our genome annotations. We first transferred these annotations to our new CI genome assembly using Mauve within Geneious Prime. We next aligned all seven genomes using Cactus 1.1 (Armstrong et al. 2019) before using the Comparative Annotation Toolkit (CAT 2.0; Fiddes et al. 2018) to transfer the annotations from the new CI genome to each of the other six genomes. Because these annotations were necessarily limited to genes that both existed and were annotated correctly in the original CI genome, we used two additional strategies to provide less biased annotations. First, we ran maker (Campbell et al. 2014; Card et al. 2019) to generate *ab initio* gene predictions for each genome, after initially training with a transcriptome generated by running StringTie (Pertea et al. 2015) on the aligned RNA-seq data and proteins taken from *D. mojavensis* and *D. melanogaster*. Second, we used PASA (Haas et al. 2003) within the funannotate pipeline (https://github.com/nextgenusfs/funannotate) to generate gene predictions after trimming, normalizing, and aligning the raw RNA-seq reads described above.

We determined *a posteriori* that the CAT annotations were by far the closest match to the raw RNA-seq data, and therefore chose to use these as our baseline for the final annotation. We next loaded GFF files from CAT, maker, and PASA, along with the raw RNA-seq alignments, into the Apollo genome annotation browser (Dunn et al. 2019) for manual curation. During manual curation we performed three tasks. First, we added new genes that were either unannotated in the original *D. mojavensis* genome or that the CAT pipeline did not add correctly. Second, we fixed genes that had either been incorrectly split or merged in the original annotation. Lastly, we fixed errors that were introduced due to sequencing errors in either the original Catalina Island genome or one of the six new genomes, which generally required manually fixing both the genome (see above) and the corresponding annotation.

We analyzed both the completeness of our genome assemblies and our annotations by using BUSCO (Seppey et al. 2019) to compare our own gene content against the most recent database of conserved single-copy dipteran genes (Diptera_odb10).

We generated mitochondrial annotations by transferring existing annotations from the CI mitochondria to each of the other mitochondrial assemblies using Mauve.

We used results from RepeatModeler above to calculate repeat content for each genome and BBMap stats (https://sourceforge.net/projects/bbmap/) to calculate GC content. To estimate transposable element (TE) content we used EDTA (Ou et al. 2019), which has been demonstrated to be effective in annotating non-model genomes (Bell et al. 2022). We used custom bash scripts to calculate the percentage of GC, repeats, TEs, and genes in 100kb sliding windows overlapping by 50kb, and plotted these percentages for each genome using the R package *circlize* (Gu 2014).

### Phylogenomics and divergence time estimation

We identified 12,218 single-copy orthologs across all seven genomes with OrthoFinder (Emms and Kelly 2019) using an iterative process. We first ran OrthoFinder under default parameters, separating single-copy orthologs from the remaining genes. We then re-ran the software on the remaining genes using stricter parameters, and repeating this procedure twice. In this way, we were able to capture genes that may have duplicated recently but before the common ancestor of the three species, and therefore still useful for our analyses.

We then performed codon-aware alignments of all single-copy orthologs using PRANK (Löytynoja 2014), and extracted fourfold degenerate sites from each alignment. We generated individual, unrooted gene trees using RAxML (Stamatakis 2014), and used these trees as input for consensus tree building using ASTRAL-III (Zhang et al. 2018) and MP-EST (Liu et al. 2010). All programs were run using default parameters.

After establishing a consensus tree topology, we used BPP (Flouri et al. 2018) on the entire genome and = (model 01) with 100,000 samples, a sampling frequency of 2, and a burn in of 10,000 samples, to estimate divergence times across the phylogeny. We altered the following parameters within bpp: thetaprior (3.0, 0.002) and tauprior (3.0, 0.003). All other parameters were left at default settings. Following recommendations for divergence time in *Drosophila* (Obbard et al. 2012) and previous work on *D. mojavensis* (Smith et al. 2012; Lohse et al. 2015), we used a neutral mutation rate of 3.5 × 10^−9^ (Keightley et al. 2009) and a rate of six generations per year to convert the substitution rate from BPP into age in years.

As several earlier estimates of divergence within this clade were made entirely (Reed et al. 2007) or in part (Oliveira et al. 2012) using mitochondrial data, we repeated the above analysis with the *de novo* mitochondrial genome assemblies. We first annotated thirteen known mitochondrial genes and extracted fourfold degenerate sites before running BPP model 01 using the same parameters as above for the nuclear genes. We used the mitochondrial mutation rate of 6.2 × 10^−8^ per site per generation (Haag-Liautard et al. 2008) and a rate of six generations per year to calculate the BPP estimate of divergence in years.

### Analysis of structural genome evolution

We aligned all seven genomes using NUCmer in order to identify breakpoints and visualize previously identified chromosomal inversions on Muller elements A, B, and E. We made figures of genome wide synteny using Dot (https://github.com/marianattestad/dot). Prior to analyzing structural variation quantitatively, we used these breakpoints to manually create “uninverted” chromosomes, wherein we forced all chromosomes to be homokaryotypic with CI. This allowed us to compare synteny inside and outside of major inversions in an unbiased manner. We re-ran NUCmer on the “uninverted” genome assemblies and used this output as input for identification of structural variation and syntenic genome regions using SyRI (Goel et al. 2019). Using the CI genome as our template, we followed Jiao and Schneeberger (2020) in quantifying the percentage of syntenic sequence in 100kb regions of the genome over 50kb sliding windows using custom bash scripts. We compared synteny across chromosomes within each genome using ANOVA. For Muller element F, we calculated chromosome-wide synteny after removing ∼350 kb at the centromeric end of the CI chromosome, which may be a misassembly as it has no corresponding region on any of the six *de novo* assemblies. For each chromosome with an inversion, we additionally compared the synteny outside the inversion on the centromeric end to the synteny within the inversion region using ANOVA. The region outside the inversion only included the region on the centromeric side of the inversion. We did not compare the non-inverted region on the telomeric end due to the extreme degradation of synteny near the telomere, especially in the interspecific comparisons.

### Analysis of molecular evolution

For molecular evolutionary analyses, we used the same set of aligned single-copy orthologs as used above in phylogenomic analyses, and used the best supported phylogeny from the analysis above. We analyzed dN/dS of each sequence across the entire phylogeny using Codeml (PAML; Yang 2007) with models 0, 7, and 8.

We considered two hypotheses regarding the relationship between structural and coding sequence evolution. First, we predicted that genes proximal to the inversion breakpoints would be more likely to experience positive selection. We tested this prediction by comparing the proportion of significantly positively selected genes within 1Mb on either end of a breakpoint to the rest of the genes in the genome. Second, we predicted that genes in regions of low synteny would be more likely to display signatures of positive selection. To examine this prediction, we performed linear regression to examine the relationship between the log_10_*ω* value of each gene and the synteny score between CI and NAV of the 100kb window containing the gene. We chose to display NAV as the source of syntenic data due to the fact that it displays the greatest variation in synteny while remaining correlated with structural variation in the other genomes (r_NAV-MOJ_ = 0.48, r_NAV-ARI_ = 0.73). However, we additionally performed the same analysis on the mean synteny scores of the two *D. arizonae* genomes and the three remaining *D. mojavensis* genomes to confirm this pattern. We performed all statistical analyses in R 3.6.3 (R Core Team 2020).

### Data access

All raw genomic and transcriptomic sequence data have been submitted to the NCBI BioProject database (https://www.ncbi.nlm.nih.gov/bioproject/) and are all associated with the accession number PRJNA593234. All scripts and other data are available at OSF (https://osf.io/mqvgh).

## Acknowledgments

We thank D. Kudrna for his work to produce the PacBio sequences. We thank N. Sage for assistance with genome annotations. This work was supported by the National Science Foundation (IOS-1557697 to L.M.M.). We would like to dedicate this work to Bill Heed, Marvin Wasserman and William Starmer whose foundational work on this system has been tremendously impactful.

## Author contributions

K.M.B., C.W.A., C.C.J., and L.M.M. conceived and designed the study. C.W.A. and C.C.J. assembled genomes. K.M.B., C.W.A., F.D., X.C., and L.M.M. annotated genomes. K.M.B., C.W.A., and L.M.M. performed analyses of genome structure. K.M.B. and M.J.S. performed analyses of phylogenomics and divergence time estimation. K.M.B., C.W.A., and L.M.M. performed molecular evolutionary analyses. K.M.B. and L.M.M. wrote the paper with input from all authors.

