## Supplemental Figure S1 for "Chromosome-length genome assemblies of cactophilic *Drosophila* illuminate links between structural and sequence evolution"

Element B (chromosome 3)

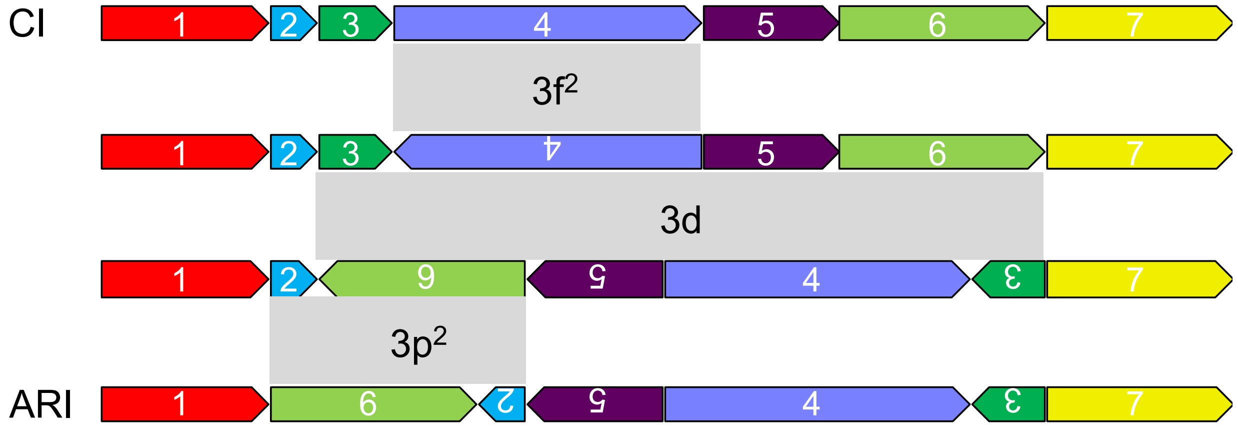


Element B (chromosome 3)

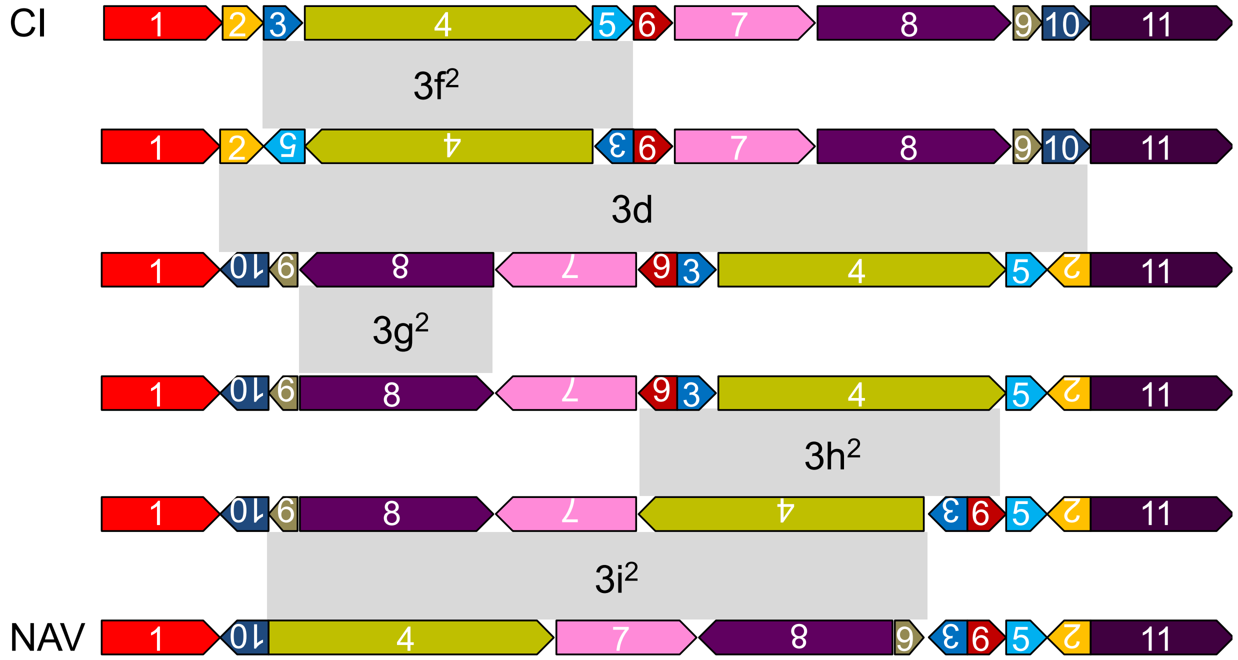

Element B (chromosome 3)

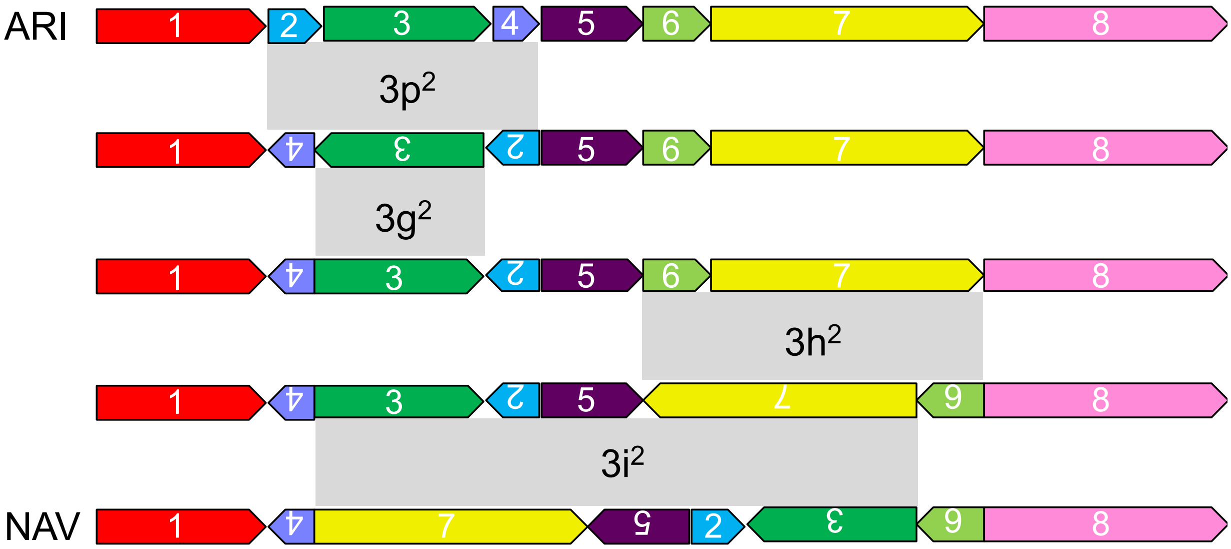


Element E (chromosome 2)

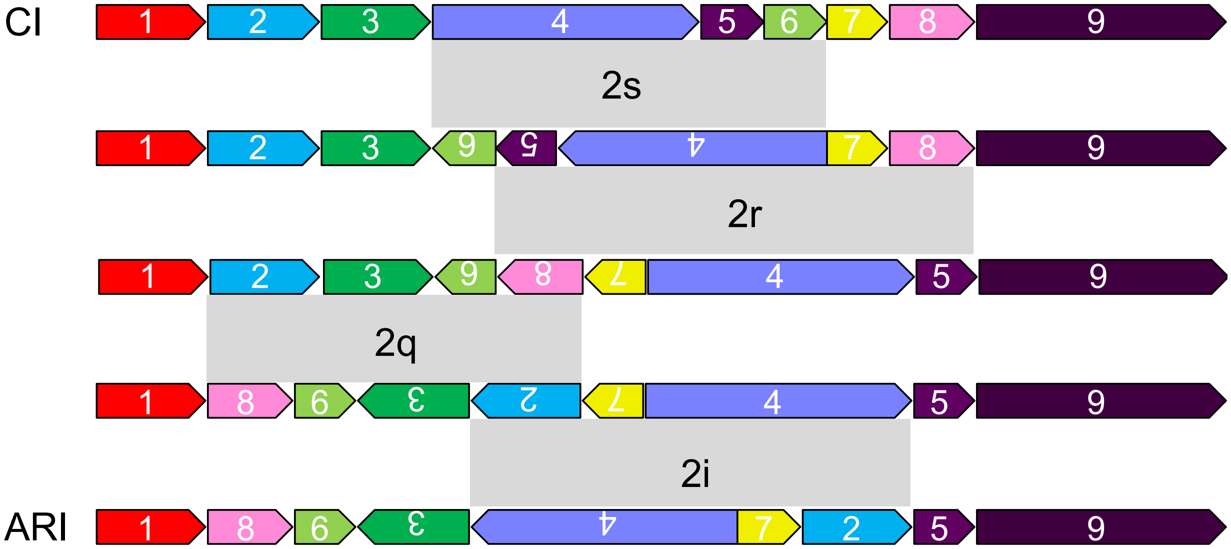

Element E (chromosome 2)

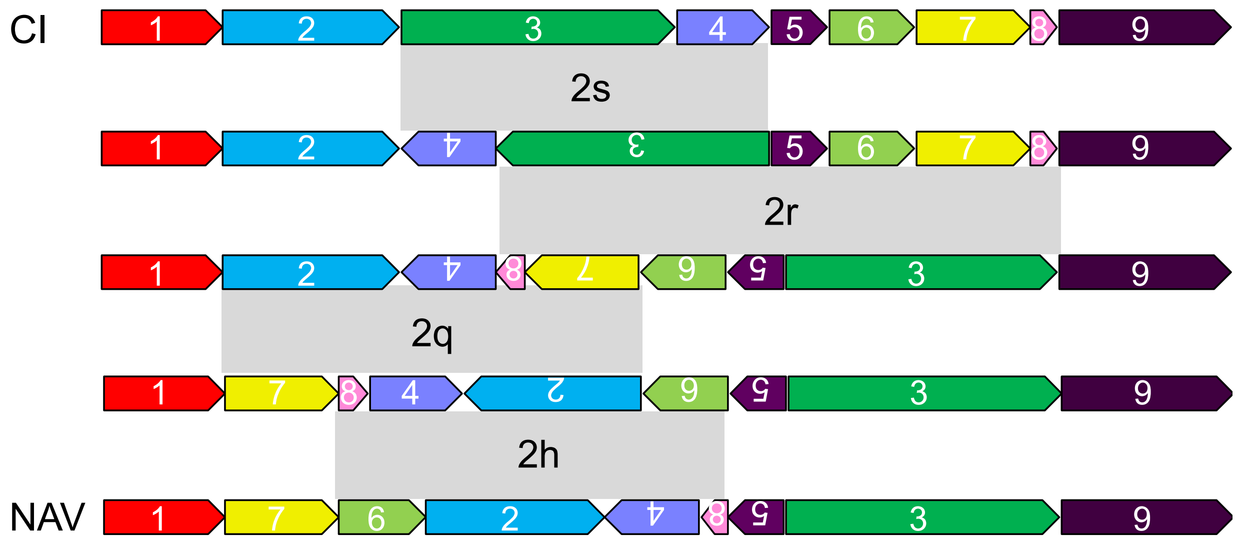


Element E (chromosome 2)

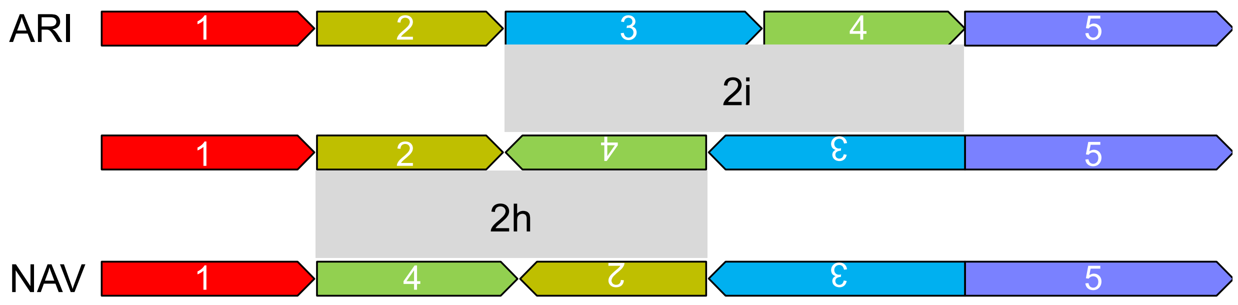


**Supplemental Figure S1.** Steps leading to the current karyotypes of the *D. mojavensis* Catalina Island, *D. arizonae* Sonora, and *D. navojoa* genomes.
