## Supplemental Figure S2 for "Chromosome-length genome assemblies of cactophilic *Drosophila* illuminate links between structural and sequence evolution"

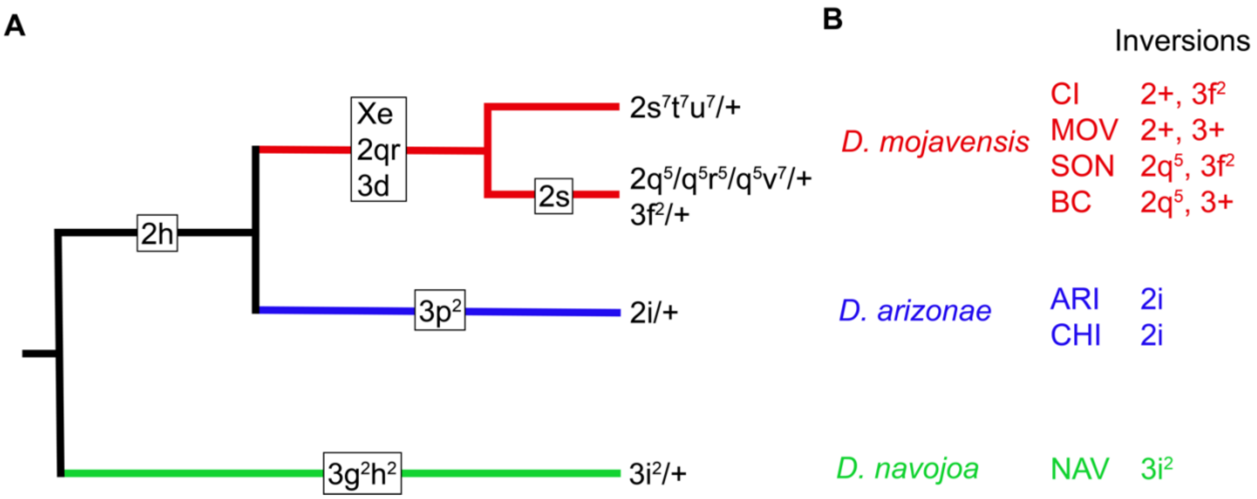


**Supplemental Fig S2. A.** Evolutionary history of chromosomal inversions within the *mojavensis* cluster as reviewed in and using the same nomenclature as Ruiz et al. 1990. **B.** Karyotype of all the genome lines used. Numbers indicate chromosome whereas muller elements A-F are chromosomes X, 3, 5, 4, 2 and 6, respectively.
