## Supplemental Figure S3 for "Chromosome-length genome assemblies of cactophilic *Drosophila* illuminate links between structural and sequence evolution"

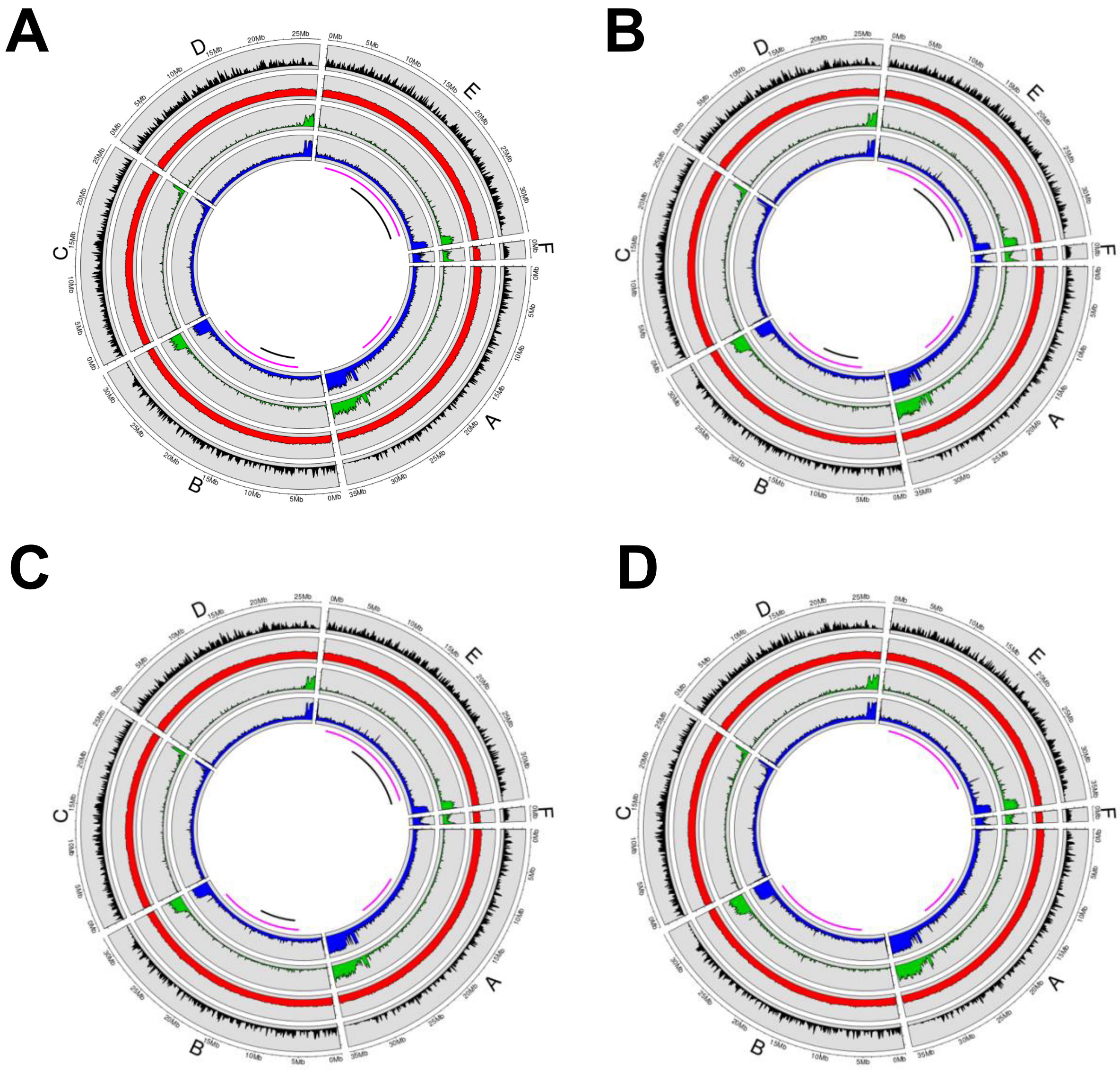


**Supplemental Figure S3.** Genome statistics for: (*A*) Catalina Island *D. mojavensis*; (*B*) Baja California *D. mojavensis*; (*C*) Sonora *D. mojavensis*; (*D*) Sonora *D. arizonae*. From outside to in, circles represent gene content, GC content, TE content, and total repeat content. Pink bars below the circles represent the regions covered by interspecific inversion polymorphisms, and black bars represent regions covered by inversion polymorphisms within D. mojavensis.
