## Supplemental Figure S4 for "Chromosome-length genome assemblies of cactophilic *Drosophila* illuminate links between structural and sequence evolution"

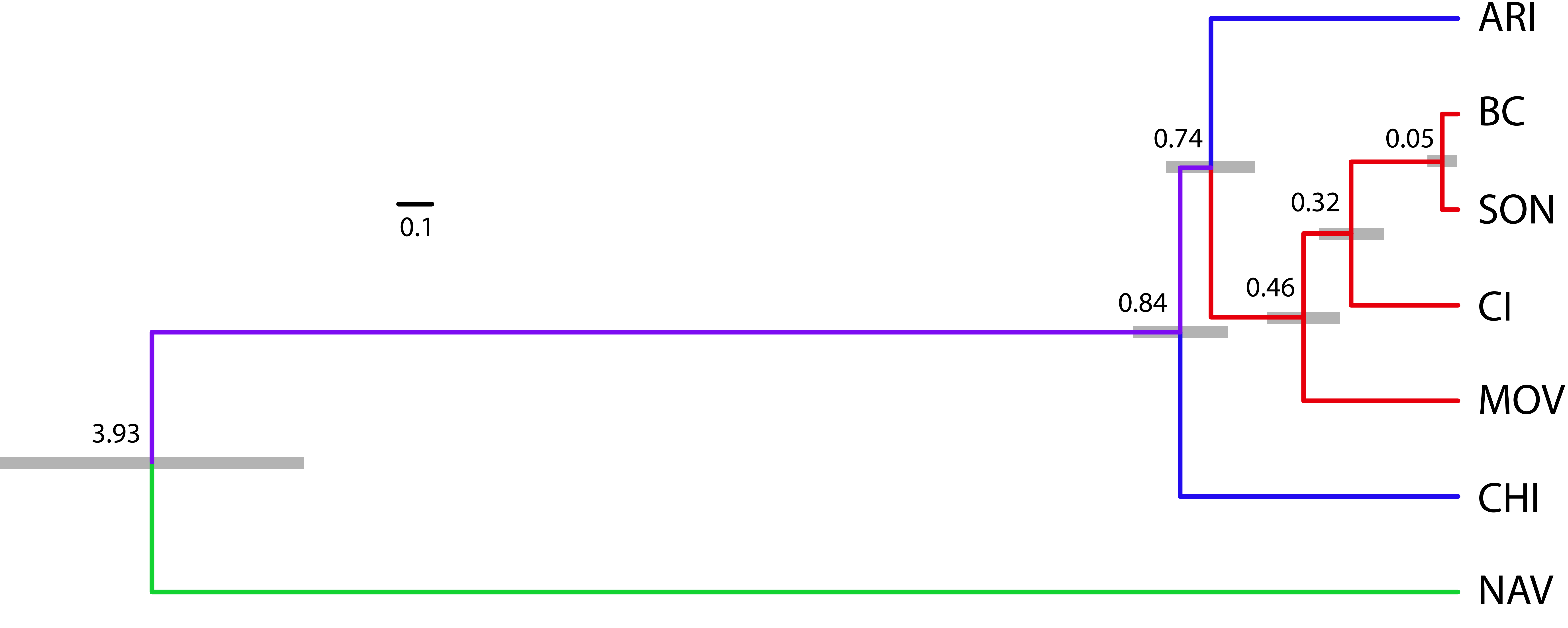


**Supplemental Figure S4.** Phylogeny and divergence times (mya) as estimated by 12,218 single copy nuclear genes. Colors represent the accepted species identities and grey bars represent 95% confidence intervals for divergence time estimates.
