## Supplemental Figure S5 for "Chromosome-length genome assemblies of cactophilic *Drosophila* illuminate links between structural and sequence evolution"

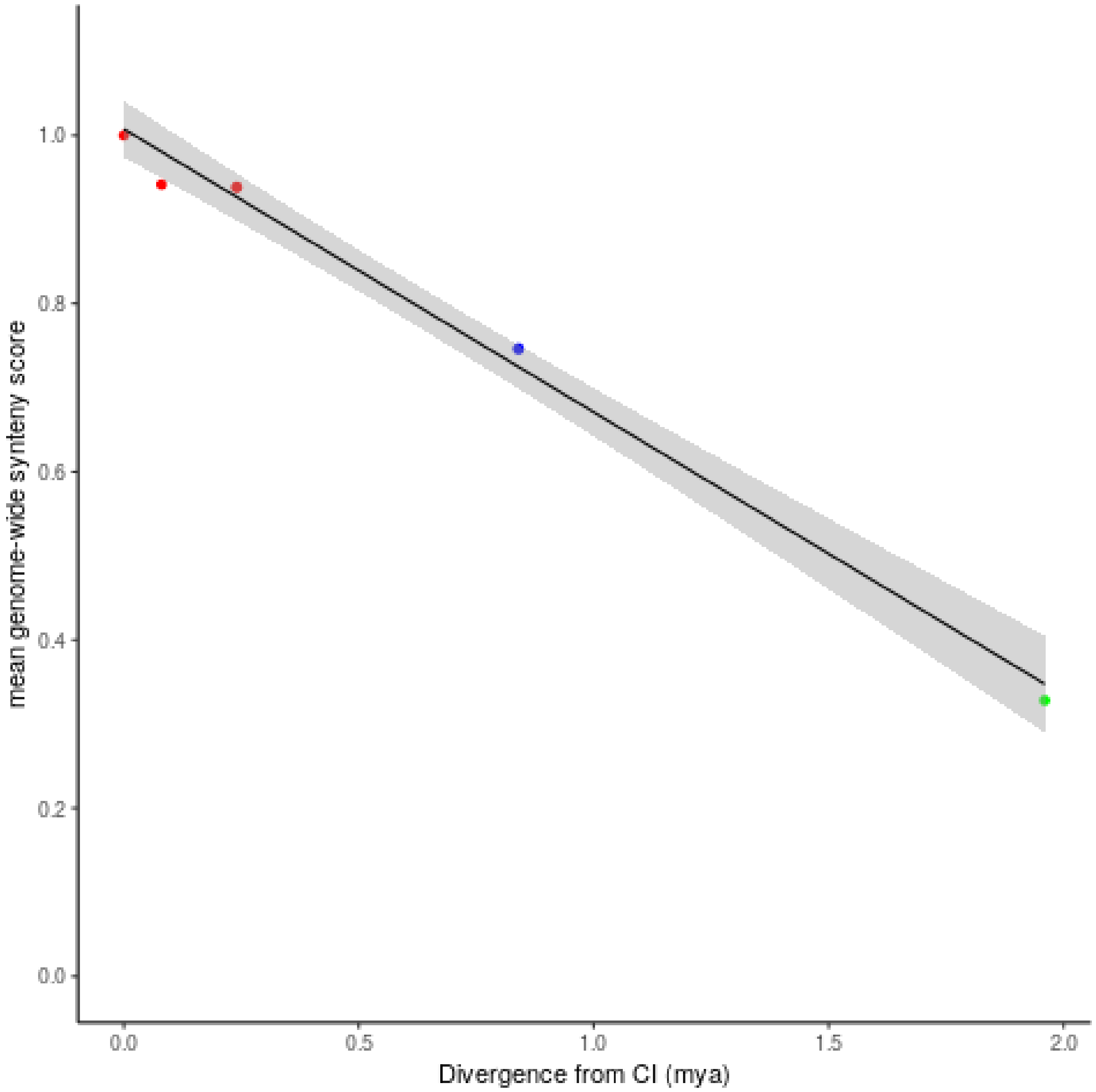


**Supplemental Figure S5.** Genome-wide synteny decay over time in the *D. mojavensis* cluster. Red points are the four *D. mojavensis* populations, blue points (nearly identical) are the *D. arizonae* populations, and the green point is *D. navojoa.*
